# Synchronous seasonal plasticity in colouration, behaviour, and visual gene expression in a wild butterfly population

**DOI:** 10.1101/2024.06.15.598356

**Authors:** Grace E. Hirzel, Noah K. Brady, Robert D. Reed, Erica L. Westerman

## Abstract

Phenotypic plasticity allows many animals to quickly respond to seasonal changes in their environment. Seasonal changes to physiological systems, such as sensory systems, may explain other more obvious changes in behaviour, often working synergistically with changes in morphology. Here we investigate if there are covarying seasonal changes to morphology, behaviour, and the visual system in the seasonally plastic butterfly *Junonia coenia*. To describe when seasonal wing patterns occur at our field sites in the central United States and for analysis of gene expression in eye tissue, we collected animals throughout the summer and fall in 2018, 2019, 2020, and 2021. For the first three years we also visited field sites to observe behaviour during focal watches and point counts throughout the flight period. We found that more *J. coenia* exhibit seasonal dark wing patterns in September and October compared to butterflies collected in previous months. This change in wing pattern correlates to an increase in basking behaviour. Eye tissues of dark fall animals and lighter summer animals exhibit different patterns of gene expression, including clock genes and genes involved in eye pigment synthesis. Subsequent analysis of monthly variation in opsin gene expression in eye tissues confirmed that opsin genes are not differentially expressed throughout the year, though *period* gene expression is higher in the fall, and females have higher blue opsin gene expression than males. This concurrent seasonal shift in colouration, behaviour, and underlying visual physiology indicates that *J. coenia* undergoes a complex shift in phenotype that encompasses more than simple changes to thermoregulation.

## Introduction

Many animals exhibit seasonal phenotypes in response to predictable changes in their environment (West-Eberhard, 2003), a phenomenon known as adaptive seasonal plasticity. This plasticity allows animals to maintain fitness, despite experiencing a wide range of conditions or encountering conditions not experienced by their parents. By utilizing environmental cues, animals can anticipate future seasonal conditions (Bartness & Wade, 1985; Perret et al., 1998; Wikelski et al., 2000). Harsher seasonal conditions, such as those experienced during dry seasons or cold winters, might cause animals to migrate, go dormant, or show other dramatic changes in physiology and behaviour that allow them to escape or mitigate seasonal stressors (Brattström et al., 2018; Fort et al., 2013; Hondelmann & Poehling, 2007). Likewise, animals often undergo seasonal changes to facilitate breeding and raising offspring during milder times of year, in order to maximally benefit from plentiful resources (Lack, 1950; Owen-Smith & Ogutu, 2013). Consequently, the shift between living in milder versus harsh seasonal conditions requires a complex response, and seasonal phenotypes rarely differ in just one trait.

In many species, seasonal phenotypes are expressed as a suite of changes to physiology, morphology, and behaviour (reviewed in Little et al., 2020; Schroeder et al., 2020). Despite the prevalence of seasonal plasticity, however, we are still uncovering the depth of animals’ seasonal physiological responses and the integration of these responses with other seasonal traits. Changes to sensory systems, which control how animals perceive their environment, are a necessary component of their seasonal response. If changes to sensory systems mechanistically explain changes in behaviour, identifying these physiological changes may be key in understanding animals’ more overt responses to seasonal change.

Of the more obvious examples of seasonal plasticity, changes in colouration and behaviour are widely observed and serve a variety of functions. Brighter colours during the breeding season can aid animals in obtaining mates, but their conspicuousness may force animals to adapt seasonal anti-predatory behaviours (McQueen et al., 2019; Radabaugh, 1989). Some animals, such as passerine birds may return to drabber colours to migrate after the breeding season concludes (McQueen et al., 2019). In other animals, seasonal colouration serves as camouflage; for example, arctic animals lose their summer brown colouration and gain a white coat to blend into white snowy backgrounds (reviewed in Mills et al., 2018). Animal colouration may also influence thermoregulatory abilities (Jin et al., 2016; Laakso et al., 2021), as darker colouration aids in increasing body temperature at cooler temperatures (Fields & McNeil, 1988; Nice & Fordyce, 2006). Darker colouration during cooler times of year may be accompanied by reduced mating and activity levels (Kárpáti et al., 2023; McElderry, 2016). Though the ultimate causes of these seasonal colours and behaviours are understood, the proximate causes (developmental, neurological, or physiological) are less well characterized.

While seasonal changes in colouration and behaviour are described in a number of animals, seasonal changes in perception or sensory systems are not as well studied. However, an animal’s perception of its environment often influences its behaviour (Ehlman et al., 2015; Wisotsky et al., 2011) and may help explain the mechanism underlying seasonal changes in behaviour. Of the recorded instances of seasonal changes to sensory systems, most are associated with arrival of the breeding season, in species where both sexual signals and behaviour are well understood. In these seasonally breeding animals, changes in number of sensory receptors (Xie et al., 2023), sensory gene expression (Shimmura et al., 2017; Tseng et al., 2018), sensory receptor location (Hamdani et al., 2007), or sensory receptor sensitivity (Rogers et al., 2022; Sisneros & Bass, 2003) coincide with the time of year when animals are attending to signals of potential mates. These physiological changes allow animals to better discriminate and detect pertinent sexual signals, including specific auditory frequencies in calls (Rogers et al., 2022, Sisneros & Bass, 2003), chemical cues in sex pheromones (Xie et al., 2023), and colour in visual signals (Shimmura et al., 2017). The correlation between behaviour and sensory physiology strongly suggest that animals’ seasonal behaviours partially stem from seasonal changes in perception.

Seasonal phenotypes are common among butterfly species. Unlike species that experience multiple seasons throughout their lives, many butterflies only experience one; rather than observing changes within an individual, seasonal phenotypes are observed through inter-generational comparisons, called seasonal polyphenisms. Many butterfly species exhibit seasonal changes in wing colour, caused by changes in abiotic factors including humidity, temperature, photoperiod, and ultraviolet light levels experienced during development (Brakefield & Reitsma, 1991; Kato & Sano, 1987; Katoh et al., 2018). Some butterflies also exhibit developmentally induced seasonal changes in behaviour, including migratory behaviour, activity levels, strength of mate preference, and background matching (Brattström et al., 2010; McElderry, 2016; Obara et al., 2008; van Bergen & Beldade, 2019). Many butterfly behaviours listed above, as well as the ability to locate nectar sources and host plants, are mediated by colour vision (Obara & Hidaka, 1968; Pohl et al., 2011; Sauman et al., 2005; Snell-Rood & Papaj, 2009; van Bergen & Beldade, 2019). Laboratory studies on one species, the tropical butterfly *Bicyclus anynana*, found that it undergoes seasonal changes in eye size and opsin gene expression, accompanying seasonal changes in wing pattern and breeding behaviour (Brakefield & Reitsma, 1991; Everett et al., 2012; Macias-Muñoz et al., 2016; van Bergen & Beldade, 2019). Here we use the common buckeye butterfly *Junonia coenia,* found in temperate North America, to examine the covariation of several seasonal traits, including changes to the visual system.

Seasonal plasticity in *J. coenia* wing colour is well documented (Daniels et al., 2012; Scott, 1975a; Smith, 1991). This multivoltine species lives 8-10 days as an adult, breeding and emerging as an adult continuously throughout the flight season (e.g. spring through fall in temperate climates). Larvae exposed to summer-like conditions (i.e. warmer temperatures and longer photoperiod) exhibit lighter ventral wing colouration as adults while those exposed to fall-like conditions (i.e. cooler temperatures and shorter photoperiod) exhibit darker ventral wing colouration as adults (Smith, 1991). Seasonal changes in behaviour have not been as well explored, but fall morph *J. coenia* are less active than summer morphs, dispersing shorter distances and exhibiting lower activity levels (Järvi et al., 2019; Scott, 1975a). Here we take an uncommon approach to studying seasonal plasticity by observing wild populations throughout the year, allowing us to determine when specific traits appear in nature, and if the timing of these seasonal traits appears coordinated. By utilizing both field surveys and molecular approaches we uncovered potential mechanisms underlying more obvious phenotypic changes. We ask 1) when the seasonal change in wing patterns occurs in populations in northwest Arkansas, a new study area for this species located in the central prairie grasslands of the United States; 2) if seasonal changes in behaviour accompany these changes in wing colour; and 3) if seasonal changes in the visual system accompany other seasonal traits (such as changes in wing pattern or behaviour).

## Materials and Methods

### Field Sites

Survey sites included three prairie sites in Northwest Arkansas, USA (Fig S1). Woolsey Wet Prairie Restoration Area (Woolsey) is a 50 acre restored wet prairie owned by the city of Fayetteville, AR, located at 36°3’52’’N and 94°13’57’’W. Chesney Prairie Natural Area (Chesney) is an 80 acre tallgrass prairie under the control of the Arkansas Natural Heritage Commission located at 36°13’10’’N and 94°28’58’’W. Stump Prairie is a privately owned 20 acre remnant tallgrass prairie, located at 36°12’18’’N and 94°29’43’’ W. Woolsey is near the edge of Fayetteville surrounded by fallow fields slowly being developed into housing. Stump and Chesney are surrounded by hay fields and pasture.

### Morphology

To determine when *Junonia coenia* begins exhibiting fall colouration in Northwest Arkansas, we collected 2 – 9 adult *J. coenia* every other week from May – November in 2018, 2019, and 2020 at our study sites and nearby public locations. The wings of adults used for morphology were analysed in two ways. First, wings were qualitatively scored using the method outlined by Smith (1991) as tan (1), dark red (5), or an intermediate colour (2, 3, or 4). Second, we measured reflectance of wings via spectrophotometer (Ocean Insights; Jaz) to quantify wing colour. We took measurements in the area distal to where Cu2 and M3 veins met on the ventral hindwing (Fig S2). For later analysis, we compared the total area under each reflection curve. We recorded qualitative and quantitative measurements blind, in regard to both sex and collection time. We noted butterflies with heavy wing wear at the quantitative measurement location on the wing and removed these measurements prior to analysis.

### Behaviour

To investigate if *J. coenia* exhibits seasonal changes in behaviour, we conducted point counts and focal watches at our sites every other week in 2018, 2019, and 2020 on weeks we did not collect butterflies for morphology. For these surveys, a single observer recorded butterfly behaviours at Woolsey, Chesney, and Stump prairies. We visited sites between solar noon and 150 minutes after solar noon from late May to early November. During each biweekly site visit we took 5 point counts at predetermined locations (Fig S1A-C). Locations for point counts mainly stayed consistent throughout the study; however, we moved one point location at Chesney between 2018 and 2019 to capture the breadth of field site across sites, years, and focal watch locations. We recorded up to three focal watches per site visit, by recording observations via audio, then later transcribing onto spreadsheets. Focal watches lasted between 30 seconds and 5 minutes. (Those that lasted less than 30 seconds were not included in the analyses). Focal watch animals were chosen from animals counted at point count locations. In 2019 and 2020 this was expanded to include other animals observed between transects to increase sample size, with care taken not to follow the same animal twice.

We categorized behaviour as *flying*, *resting*, *basking*, *nectaring*, *courting*, *chasing*, *rejecting*, *mating*, *hovering*, *slow fluttering*, and *ovipositing*. We defined *nectaring* as butterflies feeding from flowers or spending less than one second flying to the next flower. Butterflies were categorized as *chasing* whether they were following other butterflies or being followed. While *chasing* behaviour can be part of courting in some species, for this study we defined *courting* as butterflies hovering and fluttering over another butterfly (Scott, 1975b). *Rejecting* was defined as butterflies that were fluttering their wings while being courted (Scott, 1975b). We recorded *resting* behaviour when we observed inactive butterflies that had closed wings and *basking* behaviour when we observed inactive butterflies that had open wings. We recorded *slow fluttering* behaviour when butterflies transitioned between resting and basking behaviours, with less than one second spent on either the *resting* or *basking* behaviours.

### Temperature and Daylength

We also examined how temperature and daylength affect *J. coenia* colouration and behaviour. For each week in 2018, 2019, and 2020 that we collected animals or observed behaviour, we found average weekly temperatures, average weekly temperatures for the previous week, and average amount of daylight per day for the previous week. Temperatures were obtained from records on wunderground.com that were originally recorded at the nearby Northwest Arkansas Regional Airport. We calculated the weekly average of minutes of daylight per day using NOAA’s Solar Calculations spreadsheet (https://gml.noaa.gov/grad/solcalc/calcdetails.html).

### Seasonal changes to visual system

To examine seasonal changes in visual physiology, we collected heads from wild *J. coenia* in 2019, 2020, and 2021 from our field sites. For the duration of June through October, collection usually took place once a month for a period of up to 10 days. Collection took place every four weeks, usually the first or second week of the month. The exception to this was the collection period from the end of September into October, the last collection period before weather became too cold for *J. coenia*. (The collection period that started at the end of September and lasted until the end of October will be henceforth referred to as “October.”) Due to low numbers in the field, animals were not collected in June 2021. During each collection period, we collected up to five males and five females each.

To collect heads, we caught butterflies in the period encompassing 90 minutes before and after solar noon and decapitated them within 10 minutes (though usually within 5 minutes) of capture. Heads were immediately placed in 1000 μL RNALater (Invitrogen) and placed on ice. We took collected heads back to the lab and stored them at 4° C overnight before transferring to -20° C. We dissected eye and likely some lamina (as lamina is difficult to remove from the retina of *J. coenia*, though we did our best) from heads in RNALater and homogenized tissue using mortar and pestle. We then purified RNA using Nucleospin minikits (Machery-Nagel).

### RNA-Seq

We sent 12 total RNA samples to Novogene for cDNA library prep (polyA enrichment) and paired-end 150 bp sequencing (NovaSeq PE150 on NovaSeq6000, Illumina). The samples consisted of 3 males and females from each season: Summer (June/July) and Fall (late September/October) from 2019, 2020, and 2021. We chose samples based on wing colour: summer samples came from summer animals that exhibited very light (summer morph) wings and fall samples came from animals that exhibited very dark (fall morph) wings.

We aligned mRNA sequencing reads (∼33M-70M per sample) to a *J. coenia* transcriptome (Fandino et al., 2024) and genome (*Junonia coenia* v2) on Lepbase (Challi et al., 2016) using the programs Salmon and STAR, respectively (Dobin et al., 2013; Patro et al., 2017). Pre and post alignment QC was performed with FastQC and MultiQC (Andrews, 2010; Ewels et al., 2016). All samples had over 70% of reads uniquely mapped and passed QC tests for sequence quality. We input raw counts from Salmon into DESeq2 for differential gene expression analysis (Love et al., 2014). Log2 fold change between conditions was compared with the built-in Wald test, and genes were called as differentially expressed if the Benjamin-Hochberg adjusted p-value < 0.05. We compared expression levels between multiple conditions (see Table 1).

**Table 1:**
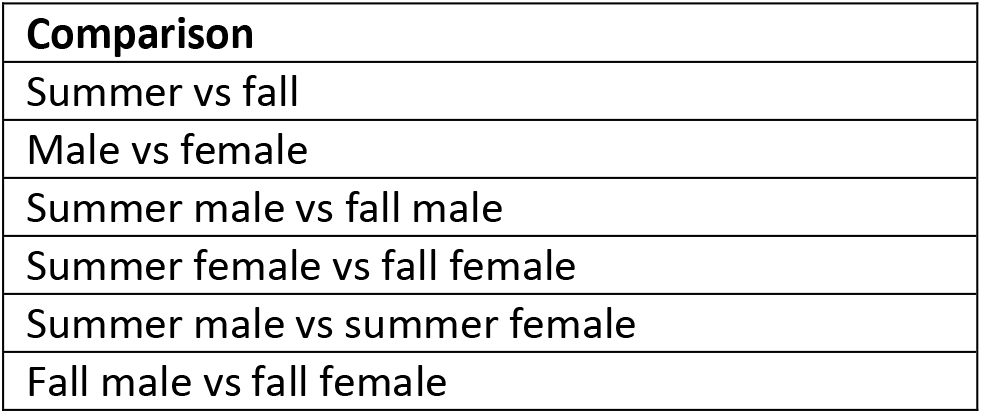
List of DESeq2 comparisons.

### qPCR

We synthesized cDNA from RNA with Oligo(dT)s following a modified protocol by Lewis and Scholes (2020), using Superscript IV with a 50 minute incubation time for reverse transcription. cDNA synthesis was immediately followed by a cleanup step using a DNA Clean & Concentrator Kit (Zymo Clean).

We then designed primers for four genes of interest – blue opsin, ultraviolet (UV) opsin, long wavelength (LW) opsin, and *period* (*per*) – and one control gene *Elongation-factor 1 alpha2* (*Ef1α-2*) using primer3.org (Koressaar & Remm, 2007; Untergasser et al., 2012). To obtain mRNA sequences we referenced the latest version of the *J. coenia* genome (*Junonia coenia* v2) found on Lepbase (Challi et al., 2016) and, for UV opsin gene, the sequenced reads from our RNA-Seq data to confirm transcripts (see Table S1 for primer sequences). We ordered primers from Integrated DNA Technologies. Before running tests, primers were tested for efficiency (See Fig S3). We ran quantitative real-time PCR (qPCR) on a CFX96 machine (Bio-Rad) with Maxima 2x SYBR Green Supermix (Bio-Rad) according to machine instructions and previously tested protocols (Lewis & Scholes, 2020). We used the resulting Ct values to find log2 fold change with the 2−ΔΔCT method (Livak & Schmittgen, 2001). We set data from the June-caught animals as our control for base comparisons, as they were the animals collected closest to the summer solstice. We included 68 males and 62 females for a total of 130 animals in our qPCR.

### Analyses

We completed subsequent analyses in R 4.2.2 (R Core Team, 2018). We tested values for normality with Shapiro tests to choose appropriate tests. To determine how collection week affected wing score we ran a generalized linear model with the glm() function and a binomial distribution, with Week 22 (May) as reference. To determine if collection week and sex influenced wing reflectance (y ∼ Week*Sex) we ran a generalized linear model with gaussian distribution and Week 22 (May) and females as reference levels. To determine how focal watch behaviour changed throughout the year we ran generalized linear models with quasipoisson distributions with Week 23 (June) acting as reference examining how the proportion of time spent on each behaviour changed with survey week. We obtained the R^2^ on figures with the *ggmisc* package (Aphalo, 2023). For point count surveys we created mosaic plots with the *vcd* package to run Pierson Chi-Square Residual tests (Zeileis et al., 2007), then created figures with ggplot2 (Wickham, 2016). To examine the correlation between wing score, basking behaviour, average temperature of collection/survey week, average temperature of previous week, and previous week’s average amount of daylength, we used the stat_poly_eq () function from the *ggmisc* package to find the values for R^2^, *p,* and the polynomial equation (Aphalo, 2023). To determine how gene expression changed throughout the year we used general linearized models to see how interactions between month and sex affected log2 fold change, with “October” and females used as the reference levels.

### Ethical Statement

Permission to survey at Stump was granted by Ozark Ecological Restoration, Inc. and permission to survey at Woolsey was provided by Environmental Consulting Operations, Inc. Animals were collected with the following permits through the Arkansas Game & Fish Commission: 40620181, 42220192, 050820201, and 072120213. We surveyed and collected at Chesney with permission from the Arkansas Natural Heritage Commission under research permits S-NHCC-18-023, S-NHCC-19-015, and S-NHCC-20-010, and S-NHCC-21-018.

## Results

### Wings begin showing fall-like characteristics in September and October

We found qualitative wing score depends on week collected (*p* < 0.001, N = 177). Wings are significantly more likely to show fall (dark) morph in late September compared to spring collected animals (*p* < 0. 017, Fig 1A). Quantitative appearance of wings, wing reflectance, also varies with collection week, (*p* < 0.001, N = 166) with the likelihood of darker wings significantly increasing in the beginning of October compared to spring collected animals (*p* < 0.015, Fig 1B).

**Fig. 1:**
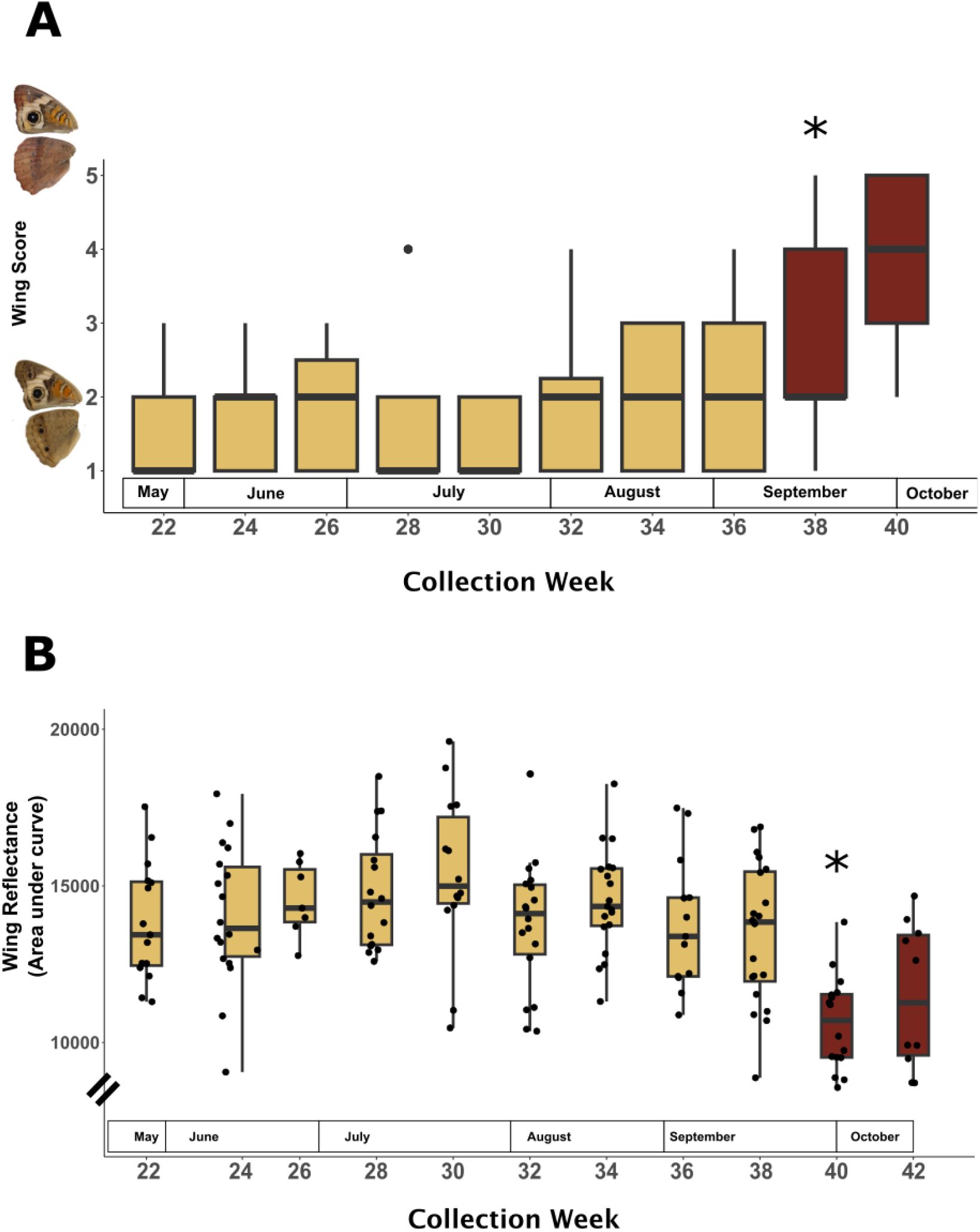
Seasonal changes in wing colour in terms of qualitative (A) and quantitative (B) characteristics. In both qualitative and quantitative characteristics animals are significantly darker in October than in previous collection weeks. * denote significance of *p* < 0.01.

### Basking behaviour changes seasonally in both point counts and focal watches

In point counts, the number of butterflies observed doing specific behaviours changes depending on survey week (N = 133, ᵪ^2^ = 146.15, *p* < 0.001, Fig 2A). We observed more butterflies *basking* in May, September, and October. In July, we observed more butterflies *nectaring*. The numbers of butterflies observed *chasing*, *mating, courting*, and *hovering* also changes depending on survey week. The number of *flying* and *resting* butterflies does not change over the course of the season in point counts.

**Fig. 2:**
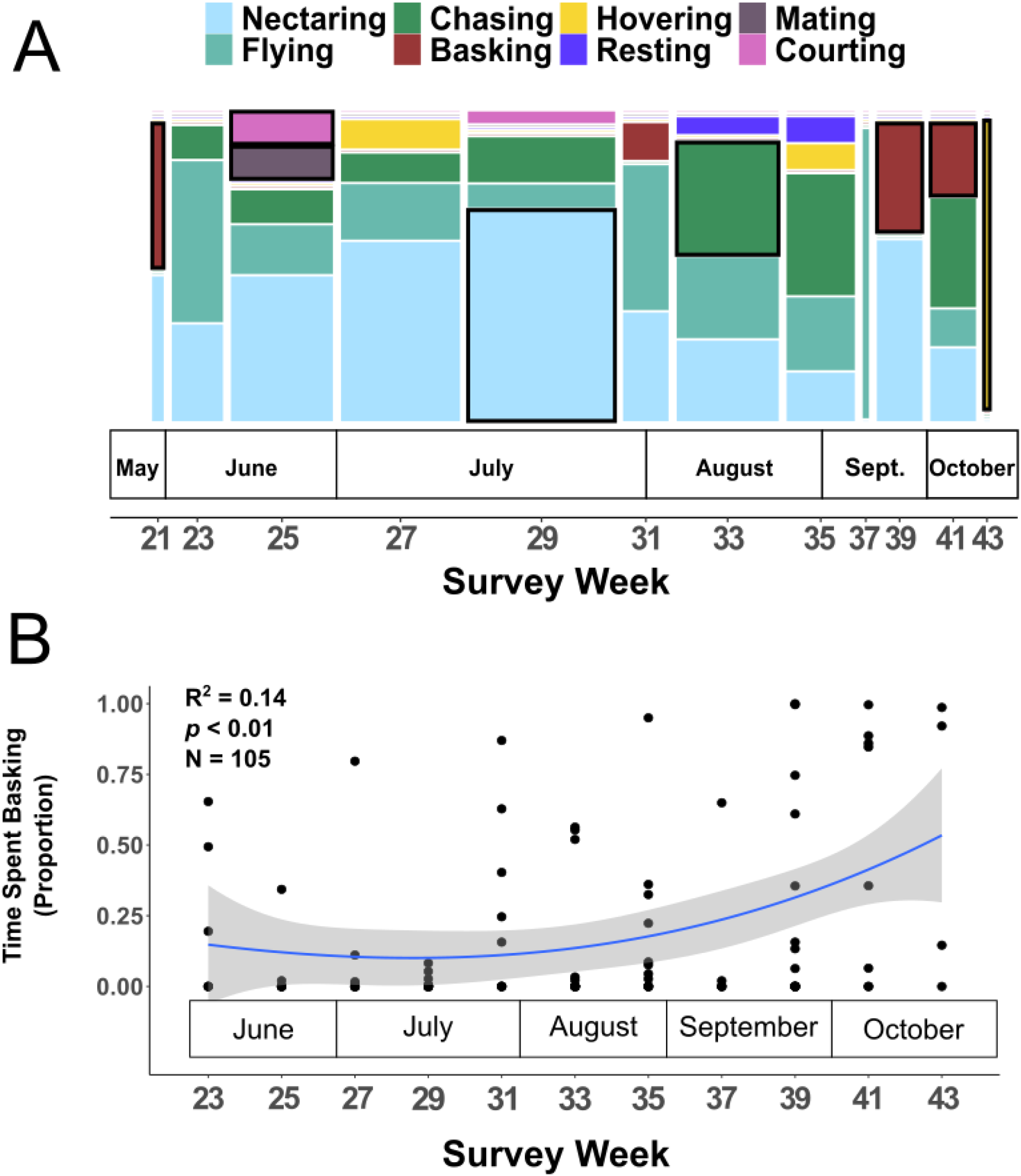
Seasonal behaviour in point counts and focal watches. Proportion of butterflies *basking*, *nectaring, chasing, mating, courting,* and *hovering* in point counts changes depending on time of year (A). Dark outlines around boxes indicate a behaviour was observed more than expected by chance. The proportion of time butterflies spent *basking* also changes throughout the year in focal watches (B).

In focal watches, the proportion of time butterflies spent *basking, flying*, and *resting* changes significantly throughout the year (N = 105, *p* < 0.01 for all three behaviours; Fig 2B, Fig S4). The proportion of time spent *nectaring* during focal watches did not change throughout the year (N = 105, *p* = 0.25; Fig S4). Of the behaviours that change seasonally, basking behaviour exhibits a trend where it increases later in the year.

### Decreases in temperature and photoperiod correlate with both increases in basking behaviour and fall-morph characteristics

The appearance of fall-like morphs corresponds to changes in daylength; as average photoperiod decreases in the week prior to collection, wings take on more fall morph characteristics (R^2^ = 0.79, *p* < 0.001, N = 28, Fig 3A) Change in qualitative wing colour also significantly correlated with temperature, increasing with decreasing previous week’s temperature (R^2^ = 0.47, *p* = 0.001, N = 28, Fig 3B). Wing colour correlates to collection week’s temperature as well, though not as strongly (R^2^ = 0.37, *p* = 0.01, N = 28, Fig S5A). The proportion of time spent *basking* similarly negatively correlates with an increase in daylength (R^2^ = 0.32, *p* = 0.012, N = 32, Fig S5B). While *basking* negatively correlates with changes in average weekly temperature of the survey week (R^2^ = 0.40, *p* < 0.01, N = 32, Fig 3C), it does not correlate with average temperatures of the previous week (R^2^ = 0.05, *p* = 0.72, N = 32, Fig S5C). Finally, proportion of time spent *basking* positively correlates with an increase in fall colouration (R^2^ = 0.32, *p* = 0.02, N =28, Fig 3D).

**Fig. 3:**
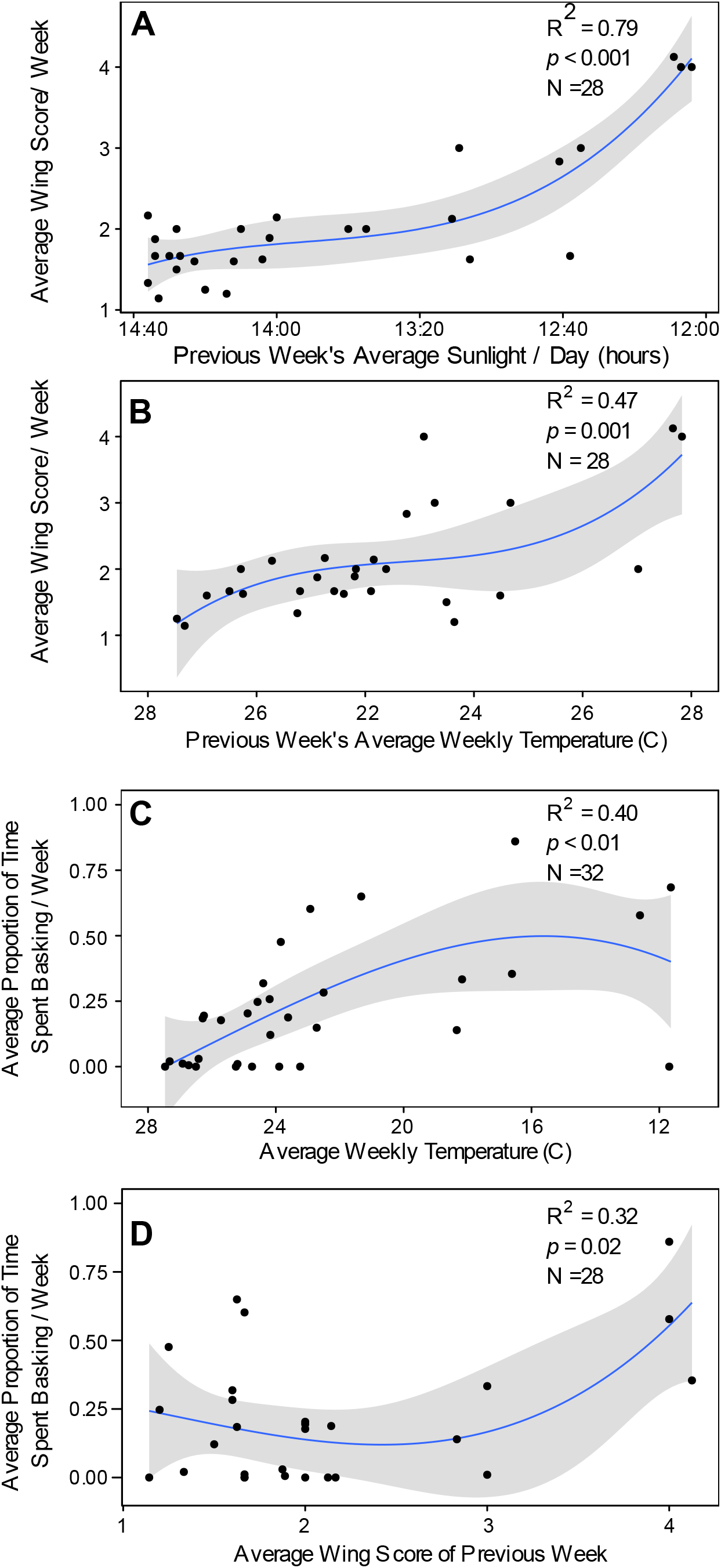
Correlations between seasonal environmental factors and butterfly traits. Wings get qualitatively more fall-like as daylength (A) and the average temperature (B) of the previous week decreases. As average weekly temperature decreases the average proportion of time spent basking during foal watches increases (C). As wings get qualitatively more fall-like the average proportion of time spent *basking* increases during focal watches (D).

### Time of year and sex affect gene expression in the eyes of *J. coenia*

Both season and sex influence gene expression in the eye. Seventy-eight genes are differentially expressed between eyes collected in summer and fall, with male and female eye tissues clustering separately in summer (Fig 4A, Table S2, Fig S6). Forty-nine genes are differentially expressed between the two sexes, with female summer and fall eye tissues clustering separately (Fig 4B, Table S3). In males, 42 genes are differentially expressed between summer and fall (Table S4). In female eyes, 27 genes are differentially expressed between summer and fall (Table S5). Among summer eye tissues, 59 genes are differentially expressed between the sexes (Table S6). Among fall eye tissues, 30 genes are differentially expressed between the sexes (Table S7).

**Fig. 4:**
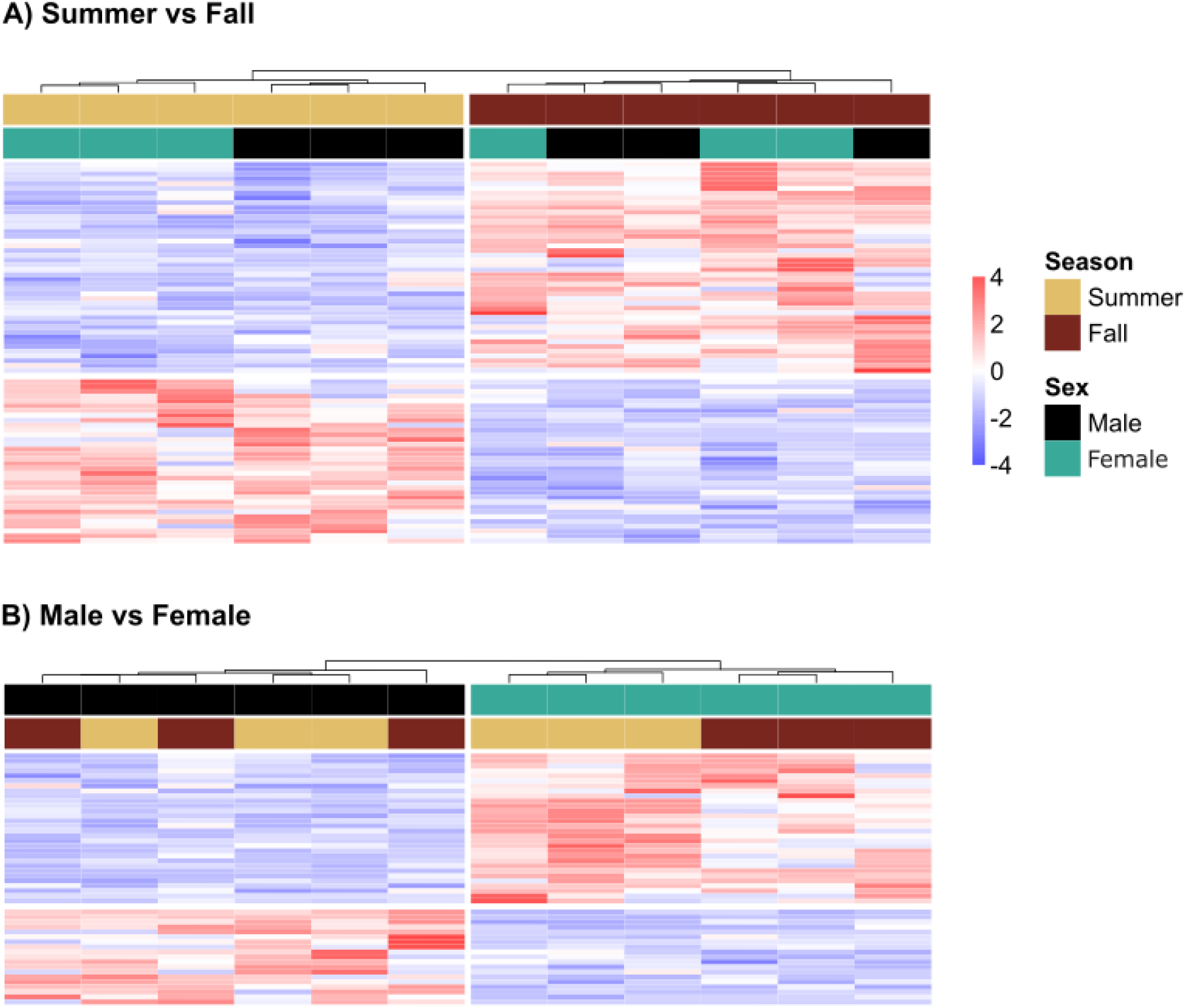
Differentially expressed genes between animals caught in summer and fall and between males and females. Gene expression in eye tissue clusters by collection season, and within summer samples, sexes cluster separately (A). Gene expression in eye tissue clusters by sex, and within females, tissues from different seasons cluster separately (B).

### Genes involved in vision and pigmentation are seasonally expressed in *J. coenia* eyes

Several genes associated with visual processing and pigmentation are differentially expressed across season and sex. Eyes express higher levels of visual genes *punch (pu)*, *vermiform (verm*), and *yellow, isoform e, (yellow-e)* in the fall compared to the summer (*pu*: p.adj < 0.01; *verm*: p.adj = 0.011, *yellow-e*: p.adj < 0.001, Fig S5, Table S2). Conversely, eye tissue expresses higher levels of the *ninaG* gene in the summer compared to the fall (p.adj = 0.023, Table S2). None of these genes are expressed at different levels in male and female eyes (*pu*: p.adj = 0.588; *verm*: p.adj = 0.810; *yellow-e*: p.adj = 0.964*; ninaG*: p.adj = 0.994), but summer male eyes express higher levels of *ninaG* than fall male eyes (p.adj < 0.01, Table S3). Fall female eyes express higher levels of *yellow-e* than summer female eyes (p.adj < 0.01, Table S5). *Cib2 (Calcium and integrin binding family member*) is more highly expressed in fall female eyes than fall male eyes (p.adj = 0.044, Table S7). Opsin genes are among the most highly expressed genes in eye tissue but are not differentially expressed between seasons or sexes (Table S8).

### Clock genes and other genes implicated in circadian rhythm differ in seasonal expression in *J. coenia* eye tissue

We also found that many differentially expressed genes are associated with maintaining circadian rhythms. Eyes of fall animals express higher expression levels of the clock genes *period* (*per*) (p.adj = 0.016) and *clockwork orange* (*cwo*) (p.adj = 0.031, Table S2). Neither gene is sexually dimorphic in expression (*per*: p.adj = 0.869; *cwo*: p.adj = 0.817) but female eye tissue expresses higher levels of *cwo* in the fall than in the summer (p.adj = 0.014, Table S5). Eye tissue expresses lower levels of *Insulin-regulator* gene *(InR*) in the fall (p.adj = 0.045, Table S2). *InR* was not sexually dimorphic in expression (p.adj = 0.326).

### Season and sex affect expression of *Doublesex*, heat shock protein, and trehalose-related genes

The expression levels of several genes involved in heat stress and sexual differentiation are affected by interactions between season and sex. Female eyes express higher levels of *Doublesex isoform F (dsx)* than males (p.adj < 0.001, Table S3), but *dsx* is also seasonal in expression in females. Summer female eyes express significantly higher levels of *dsx* compared to fall female eyes (p.adj < 0.001, Table S5). Heat shock protein genes *Hsp68* and *Hsp70ab* are not differentially expressed between sexes (*Hsp68*: p.adj = 0.674; *Hsp70ab*: p.adj = 0.820), but fall male eyes have lower expression levels of both heat shock protein genes compared to summer males (*Hsp68*: p.adj < 0.001; *Hsp70a*b: p.adj < 0.001, Table S4). Overall *Hsp68* gene expression is lower in fall animals than summer animals (p.adj = 0.018, Table S1), but *Hsp70ab* is not differentially expressed between seasons (p.adj = 0.181). *Trehelase* and *Tret1-2* (*facilitated trehalose transporter)* are more highly expressed in summer eyes (*trehelase* p.adj = 0.046, *Tret1-2* p.adj = 0.012, Table S2).

### Expression of genes associated with vision and circadian rhythm is sexually dimorphic

Female eye tissues express higher levels of genes *R and B* (*rnb)*, *dilute class unconventional myosin* (*didum)*, and *juvenile hormone acid methyltransferase (jhamt*) than male eyes (*rnb*: p.adj = 0.020; *didum*: p.adj < 0.001; *jhamt*: p.adj = 0.012, Table S3). None of these genes’ expression levels differ between seasons (*rnb*: p.adj = 0.290; *didum*: p.adj = 0.978; *jham*t: p.adj = 0.959), though *rnb* and *didum* expression is only higher in female eyes compared to male eyes in the summer and fall, respectively (*rnb* summer: p.adj = 0.017, fall: p.adj = 0.050 ; *didum* fall: p.adj = 0.049, summer: p.adj = 0.130, Table S6, Table S7) .

### Blue opsin gene expression is sexually dimorphic and *period* gene expression is seasonally expressed in eye tissue

In our more fine-scale (monthly) qPCR analysis of opsin and *period* expression patterns, we found that blue, LW, and UV opsins do not show seasonal gene expression (blue: *p* = 0.82; LW: *p* = 0.19, UV: *p* = 0.11; Fig 5A-C). LW and UV are also not sexually dimorphic in expression (LW: *p* = 0.07, UV: *p* = 0.73; Fig 5B,C). However, the blue opsin gene is differentially expressed between male and female eye tissues, with females exhibiting higher expression than males (*p* < 0.001, Fig 5A). Similar to our RNA-Seq data, *period* gene expression varies with month (*p* < 0.01, Fig 5D), and is higher in October than in previous months (*p* = 0.01). It is not sexually dimorphic in expression (*p* = 0.12; Fig 5D).

**Fig. 5:**
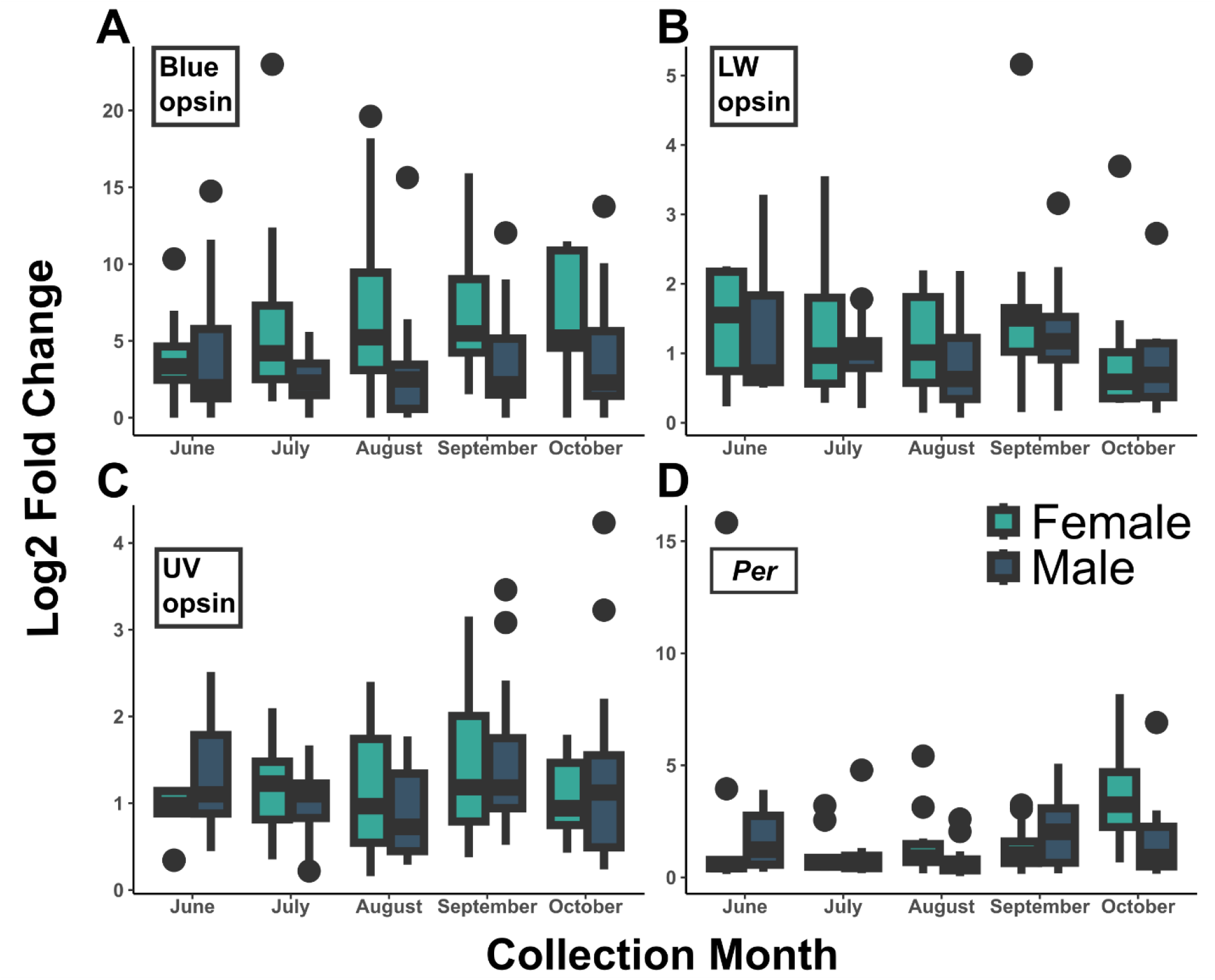
Expression of opsin and period genes throughout the year. Opsin genes show no seasonal changes in expression, but blue opsin gene is more highly expressed in female eyes (A). *Period* gene is more highly expressed in October (D).

Gene expression from one individual in July presents as the highest outlier in the blue opsin data set and is also an outlier in the long wavelength opsin and *period* data sets. Another individual in June presents as the highest outlier in the June data set for *period*. Analysis with these individuals’ data removed prior to utilizing the 2−ΔΔCT method did not cause the final results to change in degree of significance for seasonal opsin genes expression (blue: *p* = 0.45; LW: *p* = 0.19, UV: *p* = 0.10), but removing the June animal caused *period* to not exhibit significant changes in levels of expression across the year (*p* = 0.50).

## Discussion

We found that *J. coenia* exhibits a complex seasonal response as observed in covarying changes to colouration, behaviour, and visual physiology. In Northwest Arkansas populations, the dominant colour pattern of *J. coenia* shifts from the summer morph to the fall morph in September. The appearance of this dark morph correlates to increased basking behaviour, both in terms of numbers of *J. coenia* butterflies observed *basking* and proportion of time a butterfly spent *basking*. Furthermore, the eye tissue from dark winged butterflies caught in the fall have a suite of differentially expressed genes, including phototransduction genes, when compared to eye tissue from lighter summer butterflies. Our results indicate that *J. coenia* is therefore one of the growing number of known species where seasonal colouration covaries with other seasonal traits, including changes in sensory physiology and behaviour.

Our findings corroborate previous studies that indicate seasonal changes in colouration in *J. coenia* are part of an integrated response to survive cold temperatures (Järvi et al., 2019; Smith, 1991). In our populations, the fall-morph starts appearing later in the season than coastal populations, though assessment of coastal populations was done 30 years prior (Smith, 1991) and thus this difference may be more of an indicator of climate change than population-specific phenology. Like previously studies, we found population wide changes in wing colour correlate to seasonal changes in daylength and temperature, which act as environmental cues (Smith, 1991).

We also found that fall morph butterflies exhibit more basking behaviour, a sedentary activity used by butterflies to increase their body temperature (Kingsolver, 1985). This is similar to previous studies on populations from California and North Carolina that also found fall morph *J. coenia* differ in behaviour from summer morphs, exhibiting lower activity levels and dispersing shorter distances (Järvi et al., 2019; Scott, 1975a). The fall morph is more sedentary than summer morphs at similarly cool temperatures, despite an increased ability to thermoregulate at cold temperatures (Järvi et al., 2019). While both changes in wing colour and basking behaviour correlate to changes in previous week’s daylength, only changes in wing colour correlate to previous week’s temperature. This indicates that different environmental cues or critical periods determine seasonal colour and behaviour. Additional manipulative experiments in *J. coenia* could examine if the seasonal change in basking behaviour we observe in wild populations is also a developmentally induced trait or is from activational plasticity.

Behaviour and colouration covary in response to cooler seasonal temperatures in other nymphalid butterfly and insect species with seasonal forms, such as spotted wing drosophila (*Drosophila suzukii*) and multicoloured Asian lady beetle (*Harmonia axyridis)*. In these species, darker forms are better at surviving cooler temperatures and show more sedentary seasonal behaviours, including lower activity levels, decreased mating behaviours, and aggregating to conserve heat (Brakefield et al., 2007; McElderry, 2016; Michie et al., 2010; Shearer et al., 2016; Wang et al., 2011). Furthermore, these insect species experience a co-occurring seasonal change in physiology, such as decreased metabolism or reproductive diapause (Brakefield & Reitsma, 1991; Kárpáti et al., 2023; McElderry, 2016; Michie et al., 2010). Future studies could investigate if these changes in physiology are also part of *J. coenia’s* seasonal response.

Unlike the butterfly *Bicyclus anynana,* and many other previously studied animals with seasonal morphology (Brakefield et al. 2007, McQueen et al. 2019, Kárpáti et al. 2023), seasonal variation in breeding behaviour does not correlate with broad seasonal changes in colouration in *J. coenia*, at least in the ventral wing. In many species, including a number of butterflies, seasonal changes in colours used for sexual signalling co-occur with changes in breeding behaviours (Brakefield & Reitsma, 1991; McQueen et al., 2019; Radabaugh, 1989; Tseng et al., 2018). However, since *J. coenia* does exhibit seasonal variance in breeding behaviours that are not correlated with the switch from summer to fall morph, such as *chasing*, *mating*, and *courting*, it may be that *J. coenia* undergoes seasonal changes in sexual signals that were not part of the wing elements we investigated in our study. We examined the ventral wing surface, but future research could examine if there are seasonal changes in potential sexual signals on the dorsal wing. In *J. coenia*, ultraviolet reflectance on the dorsal wing is important for mate choice, though the importance of specific wing ornaments during courtship is unknown (Hafernik, 1982). Seasonal changes in these dorsal wing traits has not yet been investigated, but may better explain midsummer changes in breeding behaviours of *J. coenia*.

Along with differences in behaviour, fall morph and summer morph *J. coenia* have a number of significantly differentially expressed genes in their eye tissues, suggesting seasonal plasticity in visual physiology. Several vision related genes are differentially expressed, including *pu*, and *NinaG*, which in *Drosophila melanogaster* are part of the pathway responsible for screening and visual pigment synthesis in the eye (Ahmad et al., 2006; Mackay & O’Donnell, 1983). Differences in screening pigments in butterflies can explain differing colour sensitivities between sexes and species (Ogawa et al., 2013, McCulloch et al., 2022). If *pu* and *NinaG* are also involved in pigment synthesis in *J. coenia*, this species could be experiencing a seasonal change in its ability to see colour. This aspect of vision is relatively unstudied, as much of the work on seasonal colour vision has focused on changes in opsin gene expression, which is not differentially expressed in *J. coenia* between seasons. This is in contrast to another seasonally plastic butterfly, *Bicyclus anynana,* and many other visual animals in which opsins levels change seasonally and correlate to seasonal changes in breeding behaviour and colouration (Brakefield & Reitsma, 1991; Everett et al., 2012; Macias-Muñoz et al., 2016; Shimmura et al., 2017; Tseng et al., 2018). Nonetheless, gene expression patterns in *J. coenia* eye tissue clusters neatly by sex for summer but not fall, indicating that it may be a species in which sexual dimorphism in sensory physiology is greater at certain times of the year. While our data suggest that change in opsin expression is not driving seasonal changes in behaviour in *J. coenia*, other changes to the visual system could be, such as changes in visual pigments. Changes in eye size, facet number, and ommatidium anatomy could also be undergoing seasonal change and altering *J. coenia’s* perception of its environment and consequently, its behaviour.

Besides visual pigmentation genes, we found that several genes associated with seasonal plasticity are also differentially expressed across seasonal forms in *J. coenia* eyes, indicating there may be an integrated response to seasonal change and its associated temperature stress. Two heat shock protein genes associated with recovery from stressful conditions such as heat, *Hsp68* and *Hsp70ab*, (Colinet et al., 2010; Sørensen et al., 2003) are more highly expressed in the eyes of summer males. The genes *Trehelase* and *Tret1-2,* which are also implicated in surviving temperature stress (Perez & Aron, 2023; Shi et al., 2016), follow this pattern as well. Visual systems are usually not the target of heat stress studies, but the higher expression levels of these genes imply that summer animals are responding to high temperatures that fall animals are not. Interestingly, *yellow-e*, which has high gene expression in fall eye tissue, also has high expression in developing fall morph wings, where it plays a role in wing pigmentation (Daniels et al., 2014; Matsuoka & Monteiro, 2018). Future studies could examine if *yellow-e* expression is part of an integrated pathway controlling seasonal vision and colouration. Conversely, while *Trehelase* has lower gene expression in fall compared to summer in eye tissue, the inverse is true in *J. coenia* wing tissue (van der Burg et al., 2020). The lower expression of *Trehelase* and *Tret1-2* in fall eyes compared to summer eyes indicates trehalose may be playing another function in this tissue. Lastly, circadian rhythm genes *period* and *cwo* also are differentially expressed, with higher expression levels in fall eye tissue. This mirrors the pattern seen in many other insects which express higher levels of *period* under short photoperiod or cold temperatures, compared to long photoperiod or warmer temperatures (Hodková et al., 2003; Karthi & Shivakumar, 2014; Majercak et al., 1999). In these species, changes in clock gene expression cause changes in daily activity patterns, which we did not examine in *J. coenia,* but may be present.

In contrast to the RNA-Seq analysis, blue opsin gene expression is higher in females than males in qPCR. This was surprising, especially since seasonal and sex-specific patterns in UV, LW, and *period* are similar between the two analyses. Sexual dimorphism in the vision of *J. coenia* has not been described before. Compared to closely related species, its vision seems to be blue shifted (Briscoe & Bernard, 2005), but it is still unclear which specific wavelengths its opsins are most sensitive to. Sexual dimorphism in vision has been well characterized in a number of other butterfly species (Arikawa, 2005; Everett et al., 2012; McCulloch et al., 2017; Sison-Mangus et al., 2006) and is associated with different mating strategies between sexes in these systems (Everett et al., 2012; Sison-Mangus et al., 2006). Sexual dimorphism in vision could also function in other sex-specific behaviours, such as different foraging preferences or locating oviposition sites (Ehl et al., 2018; Snell-Rood & Papaj, 2009). In other butterfly and animal species, the opsin with the highest levels of expression often corresponds to those most important for mate choice (Macias-Muñoz et al., 2016; Tseng et al., 2018). Future studies could more closely examine intersexual differences in behaviours to determine what role higher blue opsin expression in females may play in the ecology of this species.

## Conclusions

Here we found that wild *J. coenia* exhibits a seasonal plastic response in many traits, with covariation in colouration, behaviour, and visual physiology. In late September, when temperatures and photoperiod decrease in central U.S. prairies, *J. coenia* is significantly darker than previous summer generations. This change in colouration coincides with an increase in basking, a sedentary, thermoregulatory behaviour. Furthermore, these dark winged, behaviourally different butterflies also exhibit differential expression of eye pigmentation, circadian rhythm, and temperature stress-associated genes when compared to lighter winged summer butterflies, suggesting they are neurogenomically distinct. This research places *J. coenia* among the small but growing number of species where behaviour, colouration, and sensory systems are observed to covary in response to seasonal conditions. Future studies could examine the prevalence of this phenomenon that affects both vertebrates and invertebrates living in seasonal environments.

## Supporting Information

Supporting documents are provided here.

## Acknowledgements

Special thank you to Joel Woolbright for the use of Stump Prairie and Pooja Panwar for help locating sites. For their help and patience in explaining qPCR and cDNA synthesis, thank you to Jeffrey Lewis and his lab, especially Tara Stuecker, Crystal Crook, and Julio Molina Pineda. Thank you to Yi Ting Ter, David Ernst, Keity Farfán-Pira, and Kiana Kasmaii for providing training and help on other molecular techniques and analyses. Thank you to Andrew D. Hirzel for providing audio-to-text dictation code for faster transferring of audio recorded focal watches to spreadsheets. Thank you to Jessica Proctor for collating temperature data. Austin Martinez provided the picture featured in Fig. S2. Thank you to Sarah Durant and Neelendra Joshi for their comments on this manuscript. This study was funded by Arkansas Game and Fish Conservation Scholarship and University of Arkansas Distinguished Doctoral Fellowship to GEH, NSF IOS 2128164 to RDR, NSF IOS 2238931and NSF IOS 1937201 to ELW, and the University of Arkansas.

## Author Contributions

GEH, RDR, and ELW conceived the idea and designed methodology; GEH collected the data; GEH and NKB analysed data. GEH and ELW led the writing of the manuscript. All authors contributed critically to the drafts and gave final approval for publication.

## Data availability statement

Colour, behaviour, and qPCR data will be publicly available as a Dryad Digital Repository XXX and RNA-Seq reads will be publicly available on the National Center for Biotechnology Information database, at XXX. Code will be publicly available on Zenodo at XXX.

## Data sources

Data obtained from wunderground.com is available in the datasets above and is cited in the manuscript.

