## Supplemental Figures and Table 1 for "Synchronous seasonal plasticity in colouration, behaviour, and visual gene expression in a wild butterfly population"

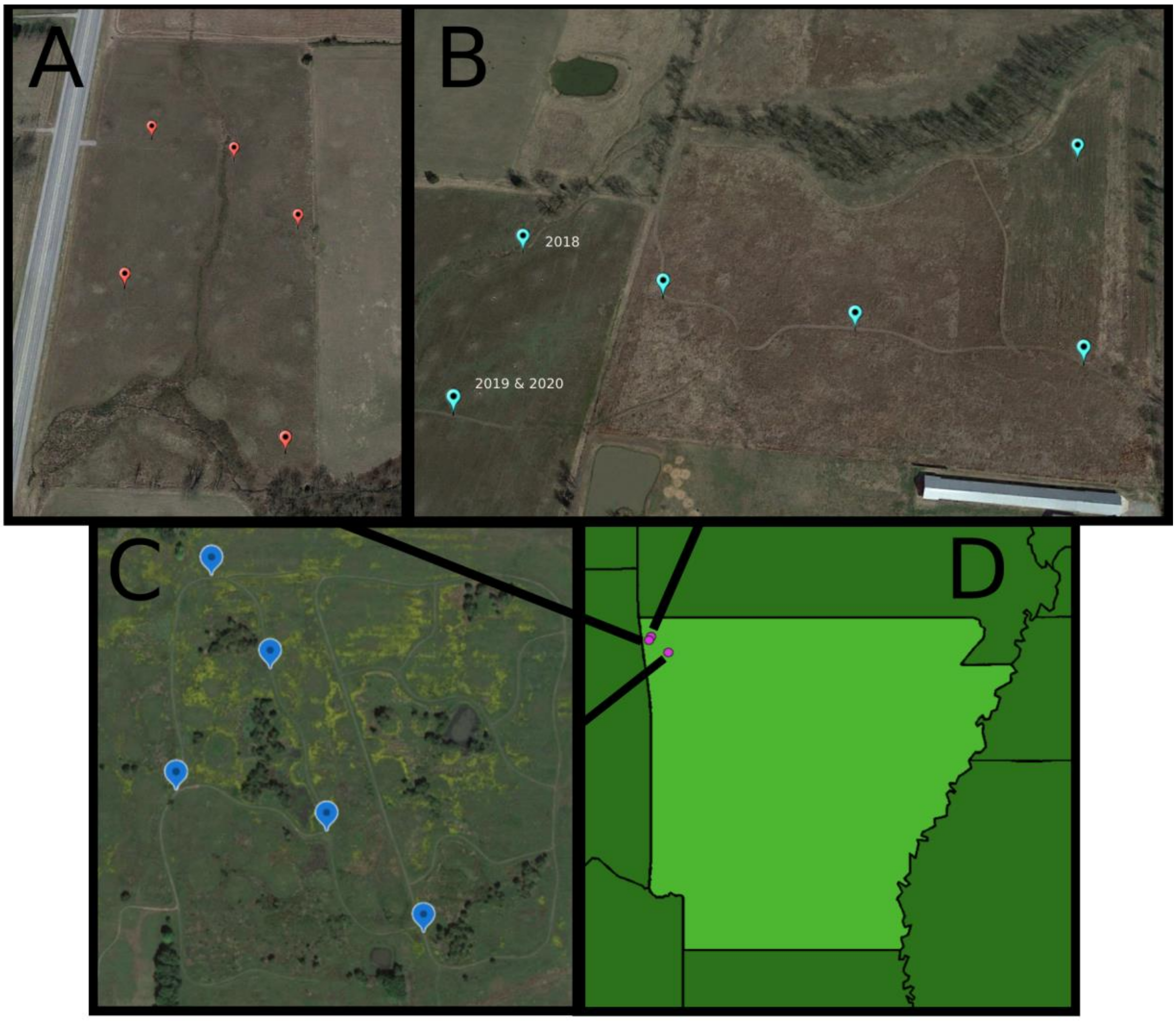


**Fig. S1: Map of field sites and locations of point counts**. Pictures are of Stump Prairie (A), Chesney Prairie (B), Woolsey Prairie (C), and map of field (D) . All point locations were consistent across years except at Chesney (B). Pictures generated using Google Earth.

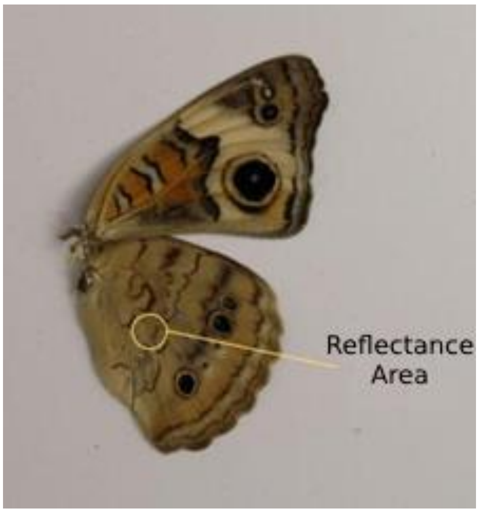


**Fig. S2: Location of reflection measurements on ventral hindwing.** Area is distal to where Cu2 and M3 veins meet. Picture used with permission.

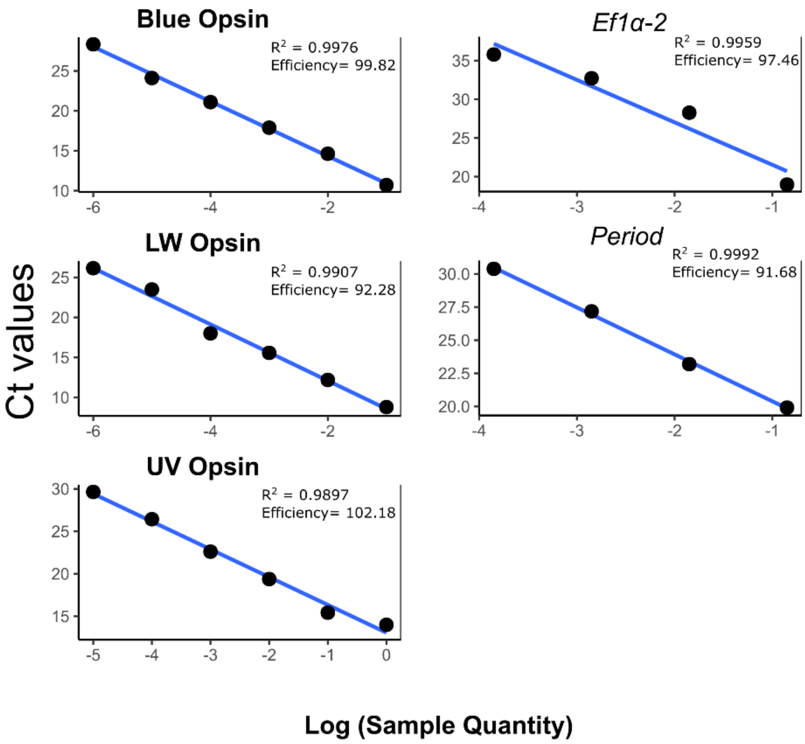


**Fig. S3: Primer efficiencies for genes of interest and control gene, *Ef1α-2*.**

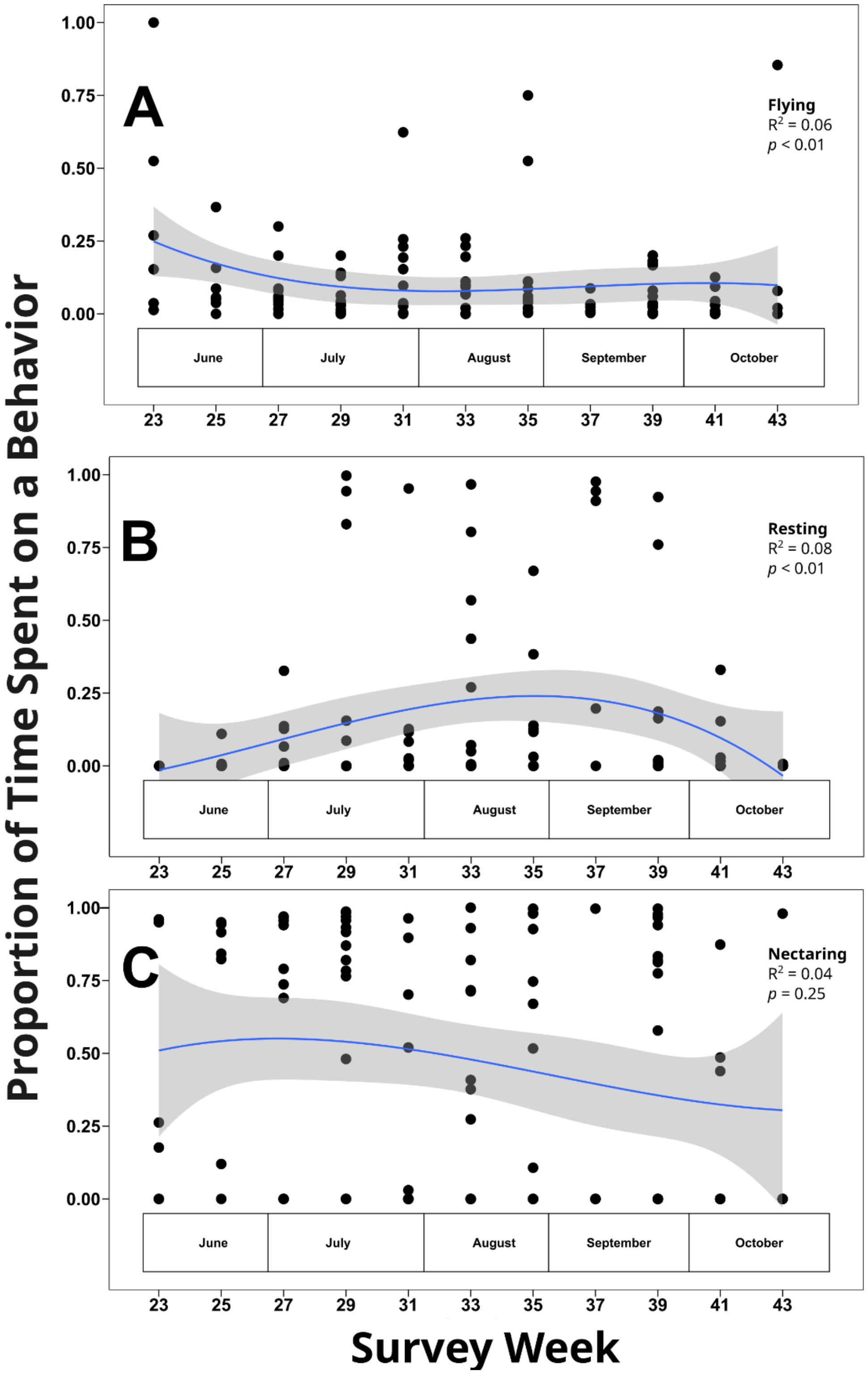


**Fig. S4: Behavior throughout the year in focal watches.** The proportion of time butterflies spend *flying* (A) and *resting* (B) changes throughout the year in focal watches. The proportion of time spent *nectaring* does not change with time of year (C).

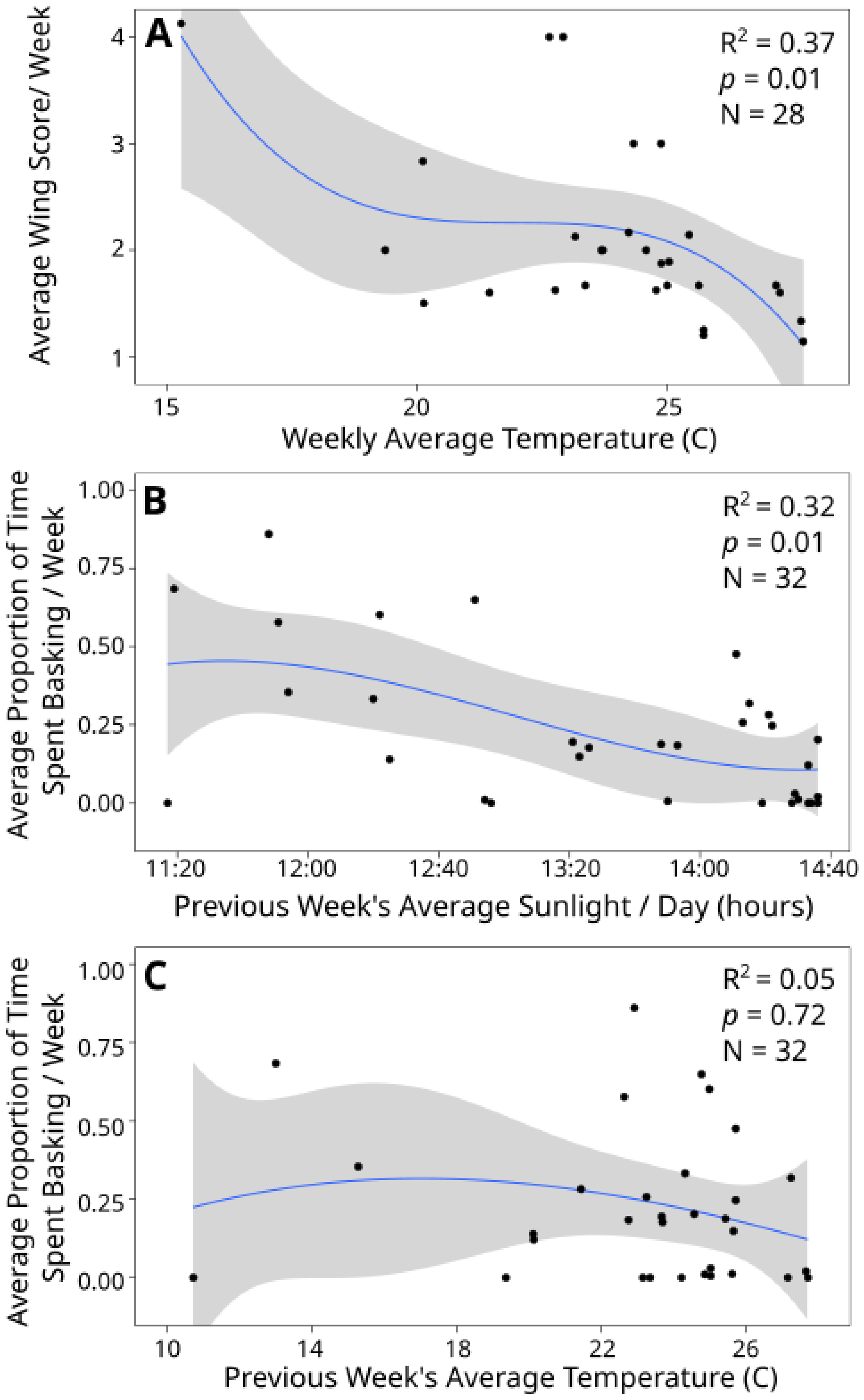


**Fig. S5: Correlations between seasonal environmental factors and butterfly traits.** As weekly average temperature decreases, wings get qualitatively more fall-like (A). As average weekly daylength increases, the average proportion of time spent basking during focal watches decreases (B). The average temperature of the week prior to survey behavior does not correlate to a change in basking behavior (C).

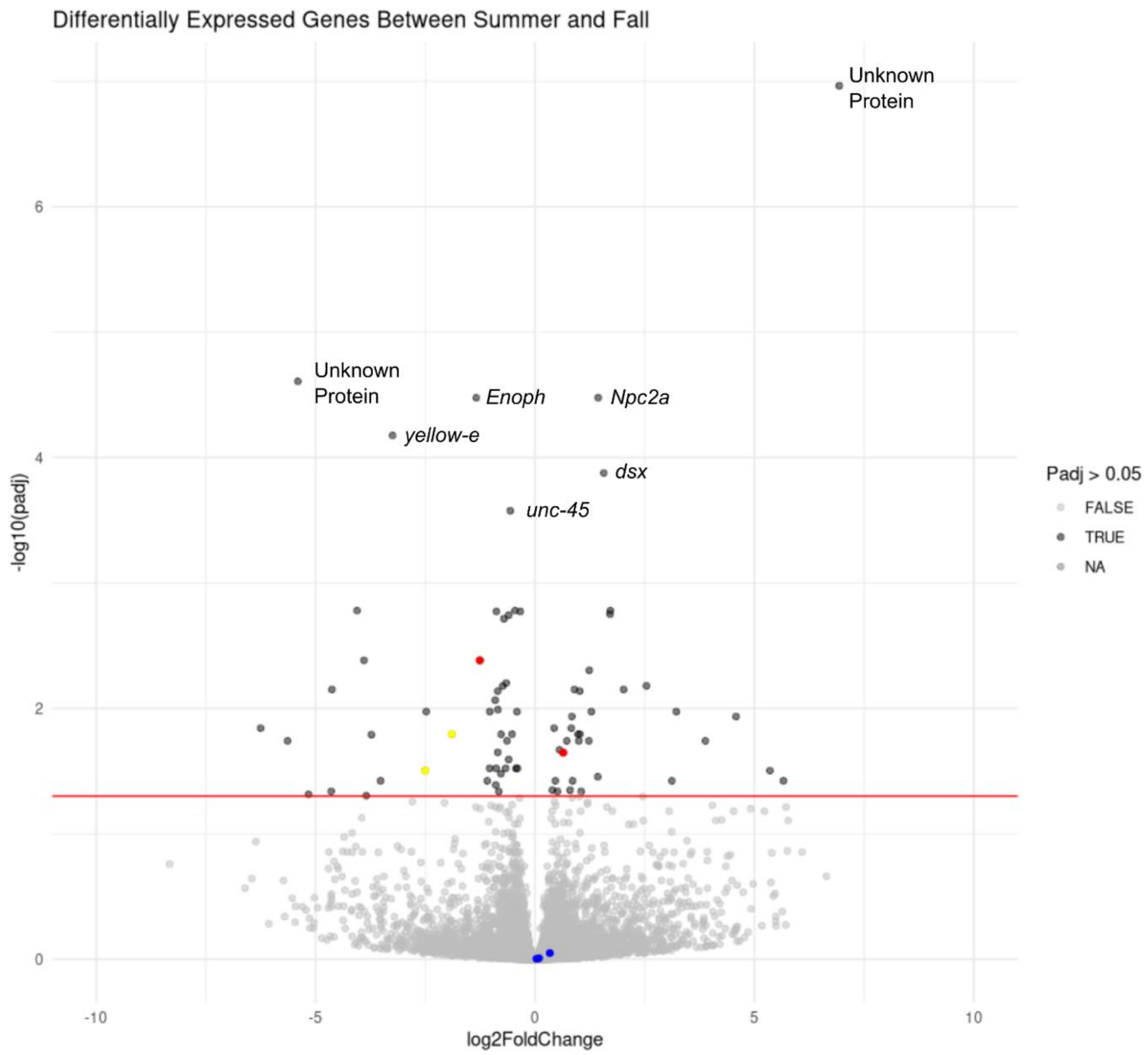


**Fig. S6: Differentially expressed genes in summer vs fall eye tissue**. Opsins are not differentially expressed, but several photopigmentation genes are upregulated in summer and some clock genes are differentially expressed. Labeled genes have a -log10(p.adj) > 3.. Blue = opsins, red = photopigmentation genes, yellow = clock genes.

**Table S1: qPCR Primer sequences for genes of interest and control gene (*Ef1α-2*)**

| Gene | Primers |
|---|---|
| Blue opsin | Forward: 5’-CGA AAC CGA CCC TTA ACG AT-3’ |
| | Reverse: 5’-TGC TCA TCA TTG CTT TTC ACG-3’ |
| LW opsin | Forward: 5’-TGC ACA CAT TTC TTA CGC CT-3’ |
| | Reverse: 5’-ACA AAG CAT CTT TCC GTC GT-3’ |
| UV opsin | Forward: 5’-CCT CGA CGG GCA CTA ATA AC-3’ |
| | Reverse: 5’-TAG GTA GTG AGG TCG CAG TC-3’ |
| *Per* | Forward: 5’-TGT TGT CAA AGT TCA CGC C-3’ |
| | Reverse: 5’-TCA CAC AAG GAA ACA CTA G-3’ |
| *Ef1α-2* | Forward: 5’-GCA AGC TAA CGA CTC AAC-3’ |
| | Reverse: 5’-ACC GAC TAC CCT CAT TTC-3’ |

**Table S2: DESeq2 results from the y ~ season model.**

**Table S3: DESeq2 results from the y ~ sex model.**

**Table S4: DESeq2 results from the y ~ season model for males only.**

**Table S5: DESeq2 results from the y ~ season model for females only.**

**Table S6: DESeq2 results from the y ~ sex model for summer only.**

**Table S7: DESeq2 results from the y ~ sex model for fall only.**

**Table S8: DESeq2 results for opsin genes.**
